# A modified decision tree improves generalization across multiple brains proteomic data sets and reveals the role of ferroptosis in Alzheimer’s disease

**DOI:** 10.1101/2024.08.07.606974

**Authors:** Mark V. Ivanov, Anna S. Kopeykina, Elizaveta M. Kazakova, Irina A. Tarasova, Zhao Sun, Valeriy I. Postoenko, Jinghua Yang, Mikhail V. Gorshkov

## Abstract

Low generalization to the patient cohort and variety of experimental conditions in the proteomic search for disease biomarkers are among the main reasons for the bumpy road of quantitative proteomics from discovery stage to clinical validation. Only a small fraction of biomarkers discovered so far by proteomic analysis reaches clinical trials. Here, we presented a machine learning-based workflow for proteomics data analysis, which partially solves some of these issues. In particular, we used a customized decision tree model, which was regulated using a newly introduced parameter, min_cohorts_leaf, that resulted in better generalization of trained models. Further, we analyzed the trend of feature importance’s curve as a function of min_cohorts_leaf parameter and found that it could be used for accurate feature selection to obtain a list of proteins with significantly improved generalization. Finally, we demonstrated that the recently introduced DirectMS1 search algorithm for protein identification and quantitation provides a simple, yet, a highly efficient solution for the problem of combining multiple data sets obtained using different experimental settings. The developed workflow was tested using five published LC-MS/MS data sets obtained in the large consortia studies of Alzheimer’s disease brain samples. The selected data sets consist of 535 files in total analyzed using label-free single-shot data-dependent or data-independent acquisitions. Using the proposed modified ExtraTrees model we found that the expressions of two proteins involved in ferroptosis Serotransferrin TRFE and DNA repair nuclease/redox regulator APEX1, are important for explaining a lack of dementia for patients with the presence of neuritic plaques and neurofibrillary tangles.

## Introduction

Biomarkers discovery is one of the important tasks for MS-based proteomics. However, one of the key issues in these studies is low generalization across data sets and/or patient cohorts.^1^

Big data analyses using machine learning (ML) algorithms bring new opportunities for the biomarker discovery by MS-based proteomics.^2^ However, this area of research is still in its infancy, undergoing a period of continuous development, leaving researchers with the best methods available at a given time. A lack of standardization in proteomic data acquisition leads to low similarity across data sets even the ones obtained for the same objects and adds further difficulties to the biomarkers discovery based on big data analysis. Note that targeted MS-based approaches widely employed for clinical trials and the diagnostic assays development based on already found protein biomarker panels do not suffer from the above issues.

Machine learning methods have multiple advantages over standard differential expression analysis based on statistical tests for biomarker candidate discovery. Firstly, ML provides a ready-to-use model for a panel of protein biomarkers. Secondly, intrinsically, these methods deal better with data containing unknown heterogeneous subgroups of patients within a group of one known pathology. Finally, the ML-based solutions can potentially uncover more complex effects of multiple protein interactions. However, there are difficulties in both detecting and interpreting these effects even upon being captured by the model.^3^

Random forest is one of the most popular choices for machine learning models in biomarker discovery applications based on computational proteomics. In this work, we proposed a modification of decision tree building in the random forest and ExtraTrees algorithm for better generalization of trained models in the context of Alzheimer’s disease proteomics. The modification allowed alternative estimation of feature importance in the trained models and better selection of a set of proteins highly relevant to Alzheimer’s disease development.

## Methods

### Data

Four proteomic data sets from large scale consortium studies of Alzheimer disease were used. The data sets are available via the AD Knowledge Portal (https://adknowledgeportal.org). The AD Knowledge Portal is a platform for accessing data, analyses, and tools generated by the Accelerating Medicines Partnership (AMP-AD) Target Discovery Program and other National Institute on Aging (NIA)-supported programs to enable open-science practices and accelerate translational learning. Data is available for general research use according to the following requirements for data access and data attribution (https://adknowledgeportal.synapse.org/Data%20Access). The following data sets were included into this study: ACT (syn5759376), Banner (syn7170616), MSSB (syn3159438) and BLSA (syn3606086). In addition, DIA data set^4^ available at ProteomeXchange^5^ under PXD025668 identifier was used. Brief description of all data sets is provided in Table 1. The data were acquired for postmortem human brain samples derived from the three types of patients: control group, Alzheimer group and Asymptomatic Alzheimer group. DIA data also divides Alzheimer’s groups into sporadic and family forms.

**Table 1.** The number of samples per condition in the data sets used in study. AD - Alzheimer group; AsymAD - Asymptomatic Alzheimer group (cognitively normal with presence of neuritic plaques and neurofibrillary tangles);

|  |  | ACT | Banner | BLSA | MSSB | DIA |
| --- | --- | --- | --- | --- | --- | --- |
| # Cases | AD | 40 | 94 | 20 | 114 | 42 |
|  | AsymAD | 14 | 58 | 13 | 21 | 11 |
|  | Control | 11 | 26 | 11 | 53 | 9 |
| brain region |  | unknown | prefrontal cortex | prefrontal cortex | frontal pole | SMTG |
| mass-spectrometer |  | Q-Exactive Plus | Q-Exactive Plus | Q-Exactive Plus | Q-Exactive Plus | Fusion Lumos |
| # of MS batches |  | 1 | 4 | 1 | 7 | 4 |
| acquisition mode |  | DDA | DDA | DDA | DDA | DIA |
| LC gradient |  | 120 min | 120 min | 120 min | 120 min | 110 min |

### LC-MS1 analysis

Data sets were analyzed using MS1-only search engine ms1searchpy (v. 2.7.3) for protein identification developed previously for ultrafast proteomics acquisition method DirectMS1^6^. Raw files were converted into mzML format using ThermoRawFileParser^7^ (v. 1.3.4). Peptide isotopic clusters in MS1 spectra were detected using biosaur2^8^ (v. 0.2.23). Parameters for the search were as follows: 5% FDR, minimum 5 scan for detected peptide isotopic cluster; minimum one visible 13C isotope; charge states from 1+ to 6+, no missed cleavage sites, carbamidomethylation of cysteine as a fixed modification and 8 ppm initial mass accuracy, and peptide length range of 7 to 30 amino acid residues. DeepLC^9^ (v. 1.1.2) software was employed for predicting retention time of peptides. Searches were performed against the Swiss-Prot human database, containing 20193 protein sequence. Decoy proteins were created by built-in ms1searchpy’s method using pseudo-shuffle method and keeping the same decoy peptides for target peptides of the same sequence from different homologous proteins.^10^

Raw to mzML file conversion (ThermoRawFileParser), peptide isotopic cluster extraction (biosaur2) and protein identification (ms1searchpy) were proceeded one-by-one for each raw file independently. At the next step, LFQ protein values were extracted within each data set group of files using directms1quantmulti script distributed along with ms1searchpy package freely available at https://github.com/markmipt/ms1searchpy.

For the quantitation, all identified peptides for a given run were sorted by intensity and only the maximum intensity was assigned to a particular combination of peptide sequence and charge state. Among the multiple peptide charge states, the one with lowest number of missing values in a data set (or the higher median intensity in case of equivalent number of missing values) was considered for subsequent analysis. All peptides with more than 50% missing values (missing value threshold) were excluded from analysis. Then, all peptide intensities were normalized by a sum of intensities for 1000 of the most intense peptides in the given run, and all missing values were replaced by run-specific minimal intensities. Then, peptide fold change was estimated by intensity value divided by the median value of this particular peptide’s intensities in the control group of the same MS batch. And finally, the protein LFQ value was estimated as the median fold change of all peptides belonging to this particular protein. All proteins with only a single peptide passing the missing value threshold were excluded from analysis.

A modified decision tree code is freely available at https://github.com/markmipt/scikit-learn under the 3-Clause BSD license. The code is a modified fork of the scikit-learn^11^ Python package (v. 1.3.1).

## Results and Discussion

### MS1 data extraction

All the data sets were analyzed using the MS/MS-free approach DirectMS1 described elsewhere^6^. There were multiple reasons for such a choice. Firstly, in our previous study it was shown that MS1-based reanalysis of MS/MS data (both DDA and DIA) can provide similar quantitation efficiency.^12^ Secondly, MS1 spectra are the most universal way to analyze and directly compare both DIA and DDA data. Finally, the MS1 spectra-based workflow provides a low number of missing values without added complexities from employing match-between-runs procedures when they are applied to the processing of large data arrays including several hundreds of files.

There were 4325, 3477, 4661, 4154, and 9646 proteins quantified for MSSB, BLSA, BANNER, ACT and DIA data sets, respectively. We kept only the intersection of quantified proteins for MSSB, BLSA, BANNER and ACT data sets, which resulted in the list of 2705 proteins. DIA data set was further used for validation of the workflow and overfitting control. Pairwise differential expression analysis for each data set, AD_vs_Control, AD_vs_AsymAD, and AsymAD_vs_Control, was added using DirectMS1Quant algorithm^10^. Differentially expressed proteins obtained in this analysis are shown in Supplementary Table S1.

### Machine learning

The ExtraTrees models^13^ based on the standard decision trees optimize mathematical objectives (such as area under curve in case of binary classification task) for the training data without regard to different groups (data sets, patients cohorts, etc.) in the data. The result is the model which provides an optimal solution on average for all samples within the training data. However, we believe that in biomedical applications, and, especially in quantitative proteomics, it is more important to obtain better generalization between data sets (cohorts) rather than just a maximal objective optimization. There are not many ways to achieve such a generalization of developed models using standard solutions. For example, one can use sophisticated weighing of samples within each data set or special feature selection procedures. Here, we propose a simple modification for decision trees, which solves the mentioned generalization issue. A new parameter, min_groups_leaf, was added to the splitting rules of decision trees. It represents the number of groups (data sets in our particular case) required to be presented in both left and right child nodes when decision tree’s split happens. A hypothetical example of the proposed method is shown in Figure 1a instead of a mathematically optimal split based on the feature “Protein10”, which results in nodes with 3 and 4 groups, the model creates a rule based on the feature “Protein2” which result in nodes with 4 and 4 groups when min_groups_leaf option is set to 4. Simply speaking, the model is forced to create rules supported by all data sets used. Indeed, such a behavior leads to a significant reduction of the overfitting, although, may suffer from the underfitting. To reduce the latter, if the parent node contains none or only one sample of some group (impossible to split into both child nodes), that group is considered as equally splitted when min_groups_leaf rule is checked. That additional behavior means that the newly introduced min_groups_leaf parameter makes more influence on the first low-depth splits and may be completely ignored at the bottom level of the decision tree. All those rules and new parameters were added into the decision tree code of scikit-learn package to modify Random Forest and ExtraTrees models. We believe that the proposed approach can potentially be added to the more complicated Gradient Boosting models like XGBoost or LightGBM.

**Figure 1.**
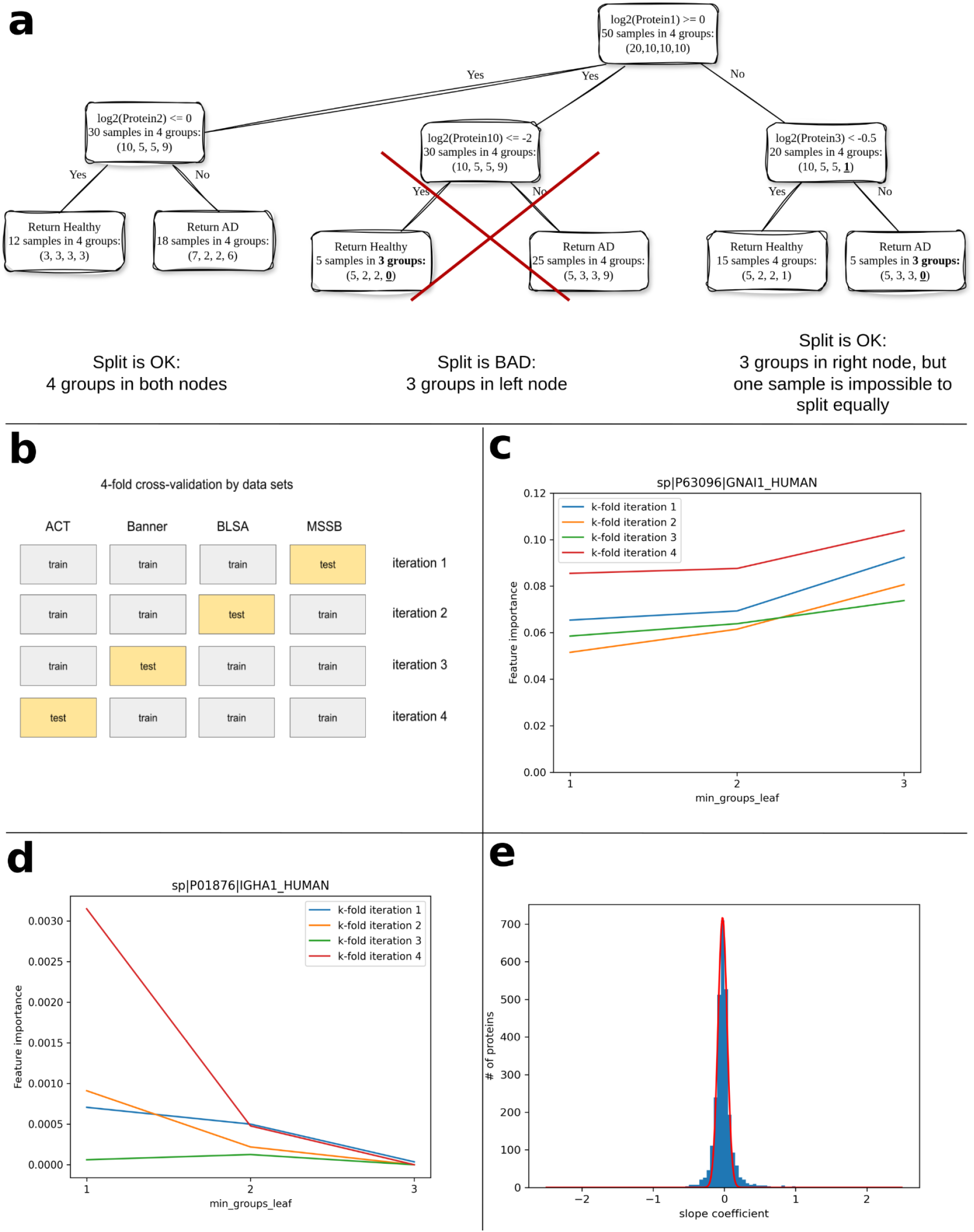
Overview of machine learning approach used in the study. (a) A visual example of proposed modification for decision trees with newly introduced parameter min_groups_leaf. (b) K-fold validation scheme with folds done by data sets. (c-d) Impurity-based feature importance for proteins GNAI1 (c) and IGHA1 (d) in each k-fold for different values of min_groups_leaf parameter for “AD vs Control” model. (e) Distribution of slope coefficients for the first k-fold iteration for “AD vs Control” model.

To test the proposed workflow we used a 4-fold cross validation scheme, in which each fold was a data set used in the study (see Figure 1b). Three different models were trained including AD vs AsymAD, AD vs Control, and AsymAD vs Control. Area under the curve (AUC) was the metric to estimate a model’s efficiency. Parameters for the models were ‘max_depth’=6, ‘min_samples_split’: 2, ‘min_samples_leaf’=1, ‘n_estimators’=500,‘max_features’=‘sqrt’, ‘n_jobs’=10. Newly introduced parameter ‘min_groups_leaf’ was varied from 1 and up to the maximal number of data sets used for training. The main results in the manuscript are shown for the ExtraTrees Regressor algorithm. Random Forest Regressor and Classifier, and ExtraTrees Classifier algorithms were also tested, yet, appeared less efficient (results are not shown). The models with maximum number of min_groups_leaf values show the highest AUC values estimated using 4-fold cross-validation (Table 2a). The DIA data used for validation have shown that the best results are achieved in two of three cases with min_groups_leaf set to 3 (Table 2b). Those results clearly demonstrate that the proposed idea provides better generalization between data sets.

**Table 2a.** Average AUC values estimated using testing splits in 4-fold cross-validation for models trained with all 2705 protein features. Bold font represents the best results.

| min_groups_leaf | Model |  |  |
| --- | --- | --- | --- |
|  | AD_vs_AsymAD | AD_vs_Control | AsymAD_vs_Control |
| 1 | 0.644 | 0.914 | 0.738 |
| 2 | 0.665 | 0.922 | 0.757 |
| 3 | **0.688** | **0.926** | **0.783** |

**Table 2b.** AUC values estimated using validation DIA data set for model trained using all 4 training data sets and all 2705 protein features. Bold font represents the best results.

| min_groups_leaf | Model |  |  |
| --- | --- | --- | --- |
|  | AD_vs_AsymAD | AD_vs_Control | AsymAD_vs_Control |
| 1 | 0.861 | 0.968 | 0.838 |
| 2 | 0.864 | 0.981 | 0.828 |
| 3 | 0.872 | **0.984** | **0.848** |
| 4 | **0.874** | 0.968 | 0.838 |

**Table 2c.**
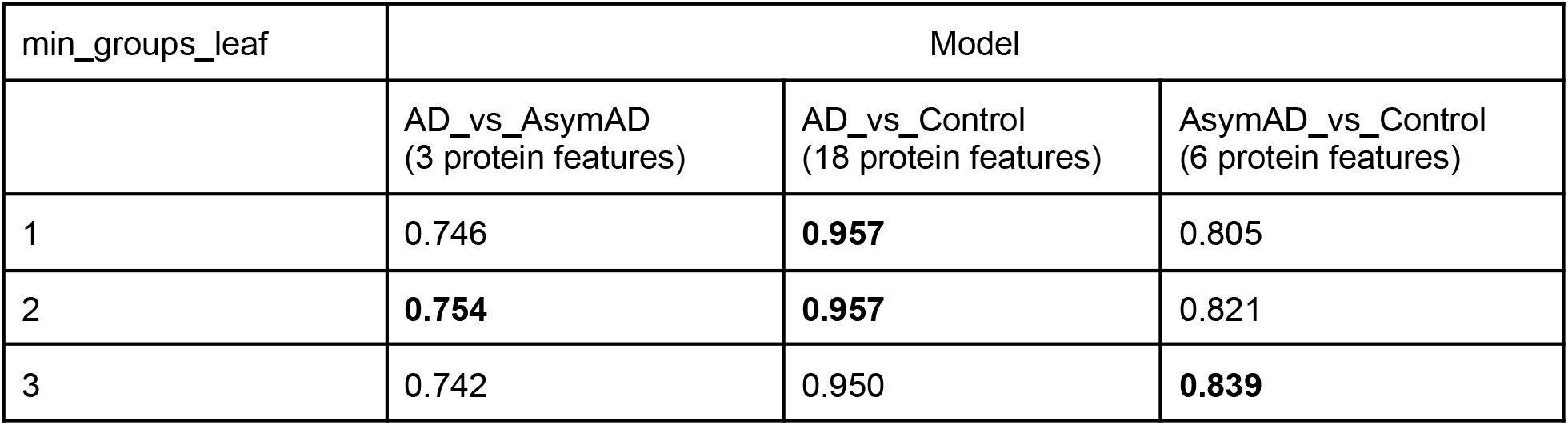
Average AUC values estimated using testing splits in 4-fold cross-validation for models trained with proposed feature selection. Bold font represents the best results.

| min_groups_leaf | Model |  |  |
| --- | --- | --- | --- |
|  | AD_vs_AsymAD<br>(3 protein features) | AD_vs_Control<br>(18 protein features) | AsymAD_vs_Control<br>(6 protein features) |
| 1 | 0.746 | **0.957** | 0.805 |
| 2 | **0.754** | **0.957** | 0.821 |
| 3 | 0.742 | 0.950 | **0.839** |

**Table 2d.** AUC values estimated using validation DIA data set for model trained using all 4 training data sets and all 2705 protein features. Bold font represents the best results.

| min_groups_leaf | Model |  |  |
| --- | --- | --- | --- |
|  | AD_vs_AsymAD<br>(3 protein features) | AD_vs_Control<br>(18 protein features) | AsymAD_vs_Control<br>(6 protein features) |
| 1 | 0.848 | **0.976** | **0.939** |
| 2 | 0.853 | 0.974 | 0.909 |
| 3 | 0.855 | 0.974 | 0.899 |
| 4 | **0.857** | 0.968 | 0.869 |

The new parameter provides an opportunity for novel feature selection procedure, which takes into account machine learning feature generalization. Within each trained model and for every protein the feature importance trend can be estimated using linear regression. The examples are shown for proteins GNAI1 and IGHA1 in Figure 1c and 1d. GNAI1 increases its importance with increasing min_groups_leaf value for all k-folds, which can be interpreted as a useful protein feature for better model generalization. On the contrary, IGHA1 decreases its importance with increasing min_groups_leaf value. The latter means that the ExtraTrees model creates more splitting rules which are not universal for all data sets when IGHA1 protein is involved. However, standard feature selection procedures based on impurity-based or permutation feature importances do not filter out such proteins. Moreover, IGHA1 is reported in the results of standard differential expression analysis for ACT data set. However, this is important when the task is generalization for big data analysis in proteomics, where acquisition schemes, experimental setups, sample preparation and data analysis are constantly evolving. To accurately estimate the feature importance for better generalization we propose probability based scoring. First, all slope coefficients within each k-fold iteration are estimated for all proteins. An example of slope coefficient distribution is shown in Figure 1e. These slope coefficients are fitted with normal distribution to obtain mean shift and standard deviation. By assuming that most proteins are not important and, thus, they provides random values for slope coefficients, the probability of chosen protein j to be randomly important in iteration i can be estimated using the following equation:

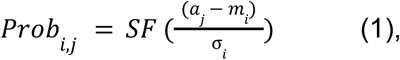

where *SF* is survival function of normal distribution, *a*_*j*_ is a slope coefficient of protein *j, m*_*i*_ and σ_i_ are mean shift and standard deviation of fitted normal distribution, respectively, and *i* is the k-fold iteration.

Then, the probability of protein to be randomly important in all iterations is:

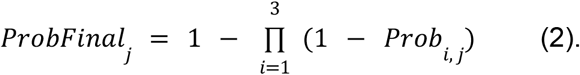

These final probabilities were calculated for all proteins and were adjusted using Benjamini-Hochberg correction. 18, 6 and 3 proteins with adjusted values of less than 0.05 were selected as important for models “AD_vs_Control”, “AsymAD_vs_Control”, and “AD_vs_AsymAD”, respectively (Table 3).

**Table 3.** List of proteins with significant p-value reported by any of five ML-based models used in the study.

| dbname | gene | p-value |  |  |  |  |
| --- | --- | --- | --- | --- | --- | --- |
|  |  | AD_vs_Control | AsymAD_vs_Control | AD_vs_AsymAD | RankedBrak | RankedCERAD |
| P63096 | GNAI1 | 0 | 0 |  | 0 |  |
| P05067 | A4 | 0 | 0 |  |  |  |
| Q9Y2J0 | RP3A | 0 |  | 0 | 1.1e-09 | 0 |
| P08754 | GNAI3 | 0 |  |  |  |  |
| Q16181 | SEPT7 | 0 |  |  |  |  |
| Q9NQC3 | RTN4 | 0 |  |  |  | 0.0055 |
| P14136 | GFAP | 2.9e-10 |  |  |  |  |
| P04792 | HSPB1 | 8.7e-08 |  | 0.00013 |  |  |
| P31146 | COR1A | 6.1e-06 |  |  |  |  |
| Q99719 | SEPT5 | 9.8e-06 |  |  |  |  |
| P60201 | MYPR | 8.7e-05 |  |  |  |  |
| P60763 | RAC3 | 0.00011 |  |  |  |  |
| Q13153 | PAK1 | 0.0026 |  |  |  |  |
| Q16623 | STX1A | 0.003 |  |  |  |  |
| Q8WYJ6 | SEPT1 | 0.0041 |  |  |  |  |
| P04899 | GNAI2 | 0.0089 | 0.014 |  |  |  |
| P02686 | MBP | 0.0094 |  |  |  |  |
| P06744 | G6PI | 0.035 |  |  |  |  |
| Q8NFI4 | F10A5 |  |  | 0.0068 |  |  |
| Q9UJZ1 | STML2 |  | 0 |  |  |  |
| P27695 | APEX1 |  | 0.0019 |  |  |  |
| P12036 | NFH |  | 0.05 |  |  |  |
| P46821 | MAP1B |  |  |  | 0 | 0.00027 |
| P61764 | STXB1 |  |  |  | 0.00011 |  |
| P02787 | TRFE |  |  |  | 0.00043 |  |
| Q5VTE0 | EF1A3 |  |  |  | 0.0052 |  |
| Q9Y2J2 | E41L3 |  |  |  |  | 0.00082 |
| Q9UPV7 | K1045 |  |  |  |  | 0.0087 |

The AUC values estimated using 4-fold cross-validation and validation DIA data set are shown in Table 2c and Table 2d, respectively. There is no optimal min_groups_leaf value for all models suggesting that this parameter is not required with small and carefully selected subsets of proteins for training.

To further analyze the proteins reported in the proposed machine learning approach, we checked if those proteins were reported in standard differential expression analysis on the same data (see Supplementary Figure S1). 9 (of 18), 5 (of 6) and 1 (of 3) of ML-obtained proteins were unique in the machine learning approach for “AD_vs_Control”, “AsymAD_vs_Control” and “AD_vs_AsymAD”, respectively. LFQ values distributions for proteins highlighted by machine approach are shown in Supplementary Figures S2-S4.

### Alternative models

Asymptomatic Alzheimer group containing cognitively normal people with presence of neuritic plaques and neurofibrillary tangles is important for studying Alzheimer disease. The classification of this group of individuals as belonging to the diseased or healthy patient cohort remains a subject of debate.^14^ This is because there is no definitive answer to the questions: whether the preclinical stage of Alzheimer’s disease will progress to clinical dementia during their lifetime, and whether this neurodegeneration is a distinguishing feature of Alzheimer’s disease or a manifestation of normal aging? It is likely that mechanisms exist preventing the development of clinical stages of Alzheimer’s disease or other neurological disorders in this group of patients. We developed special models to better reveal the insight on the mentioned topic. Instead of simple training for classification between the patient groups, we utilized Braak (neurofibrillary tangles localization) and CERAD (neuritic plaques localization) scores available with the data. Indeed, the available data is highly unbalanced. For example, 80% samples within the AD group have the highest Braak scores (5 or 6) in our data sets. On the contrary, only 19% of samples have those scores in the AsymAD group. That may result in that standard group-based model “AD_vs_AsymAD” gives a higher importance to the group of proteins which distinguish low and high Braak/CERAD scores rather than Asymptomatic vs Dementia Alzheimer patients. To deal with this problem, we used a special scheme for targets found by machine learning (Figure 2a). The idea is that the ML algorithm should produce higher training error for AD and AsymAD samples with highest Braak/CERAD values, while the samples with low Braak/CERAD values should be less important. Other parameters and training procedures were the same as described above. Using the probabilities calculated by Eq.2, we found 9 proteins reported as important by either of the considered here alternative training models, Braak- or CERAD-based: MAP1B (Braak and CERAD), RP3A (Braak and CERAD), STXB1 (Braak), GNAI1 (Braak), TRFE (Braak), EF1A3 (Braak), E41L3 (CERAD), RTN4 (CERAD) and K1045 (CERAD). The predicted values for the DIA data set using the models trained on all four Synapse’s DDA data sets are shown in Figures 2,b and c for min_groups_leaf=4 and min_groups_leaf=1 parameters, respectively. Braak-based and CERAD-based models trained with min_groups_leaf option correlate better than standard ExtraTrees model (min_groups_leaf=1). The Asymptomatic Alzheimer group was clearly divided into two subgroups close to either control or AD samples, further supporting the existing Asymptomatic Alzheimer theories. To deeper analyze detected proteins, the LFQ values grouped by Braak stage and Condition (AD, AsymAD, and Control) were shown in Figure 3. We see that most of these proteins (STXB1, RTN4, K1045, GNAI1, and RP3A) decrease in concentration for both AD and AsymAD groups with the increase in Braak stage. We believe that these proteins do not really differentiate AD and AsymAD groups. Protein EF1A3 has a more interesting pattern: it increases back to Control values at Braak stage 6 for both AD and AsymAD groups. Proteins E41L3 and MAP1B both decrease in AD group, but remain stable in Control and AsymAD group (except Braak 6 stage). The remaining protein TRFE decreases in AD, remains stable in Control and increases in AsymAD group (except Braak 6 stage)

**Figure 2.**
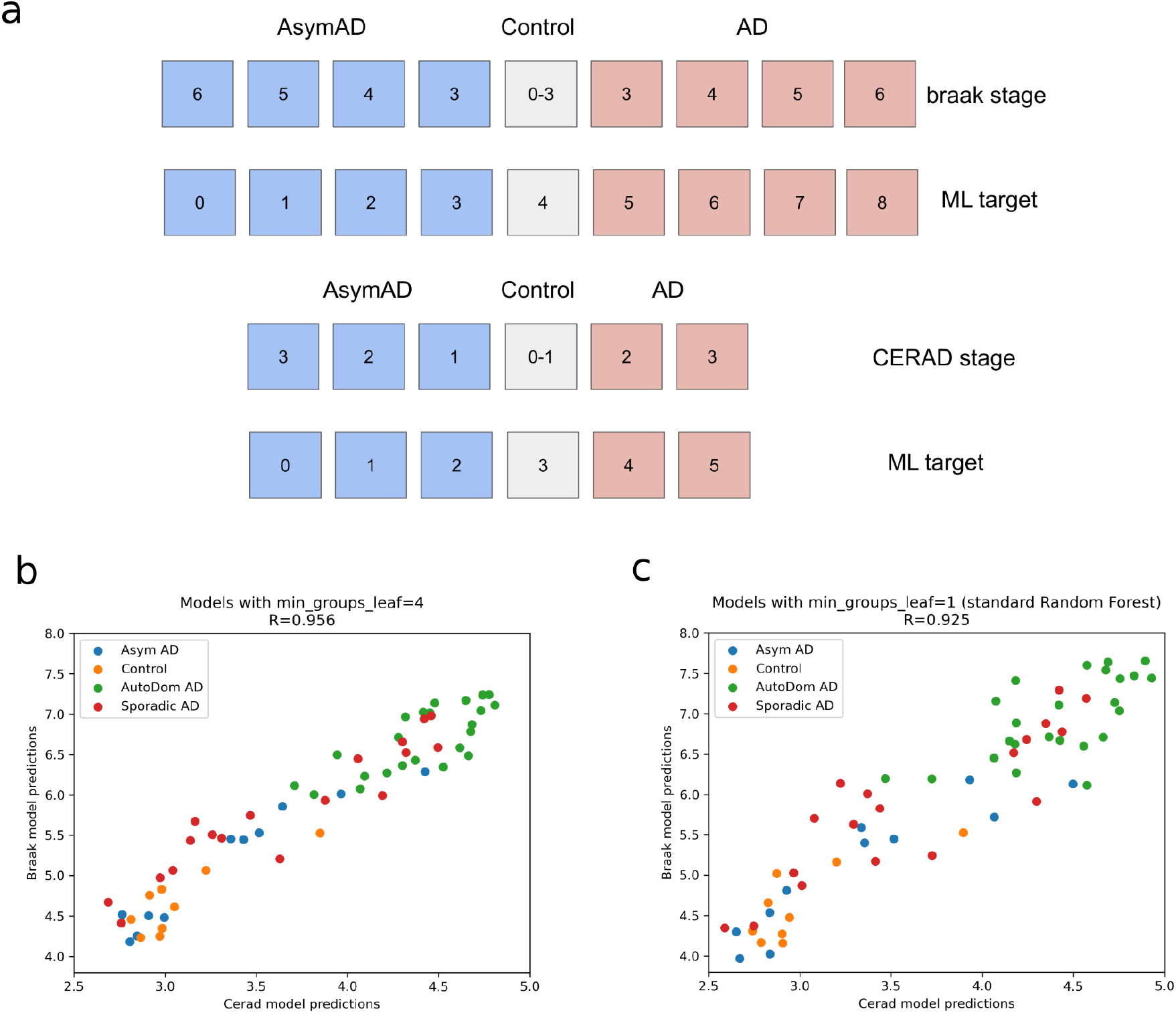
(a) Custom targets for machine learning based on braak and CERAD stages; Braak-based and CERAD-based model predictions for DIA data samples using min_groups_leaf=4 (b) and min_groups_leaf=1 (c) options.

**Figure 3.**
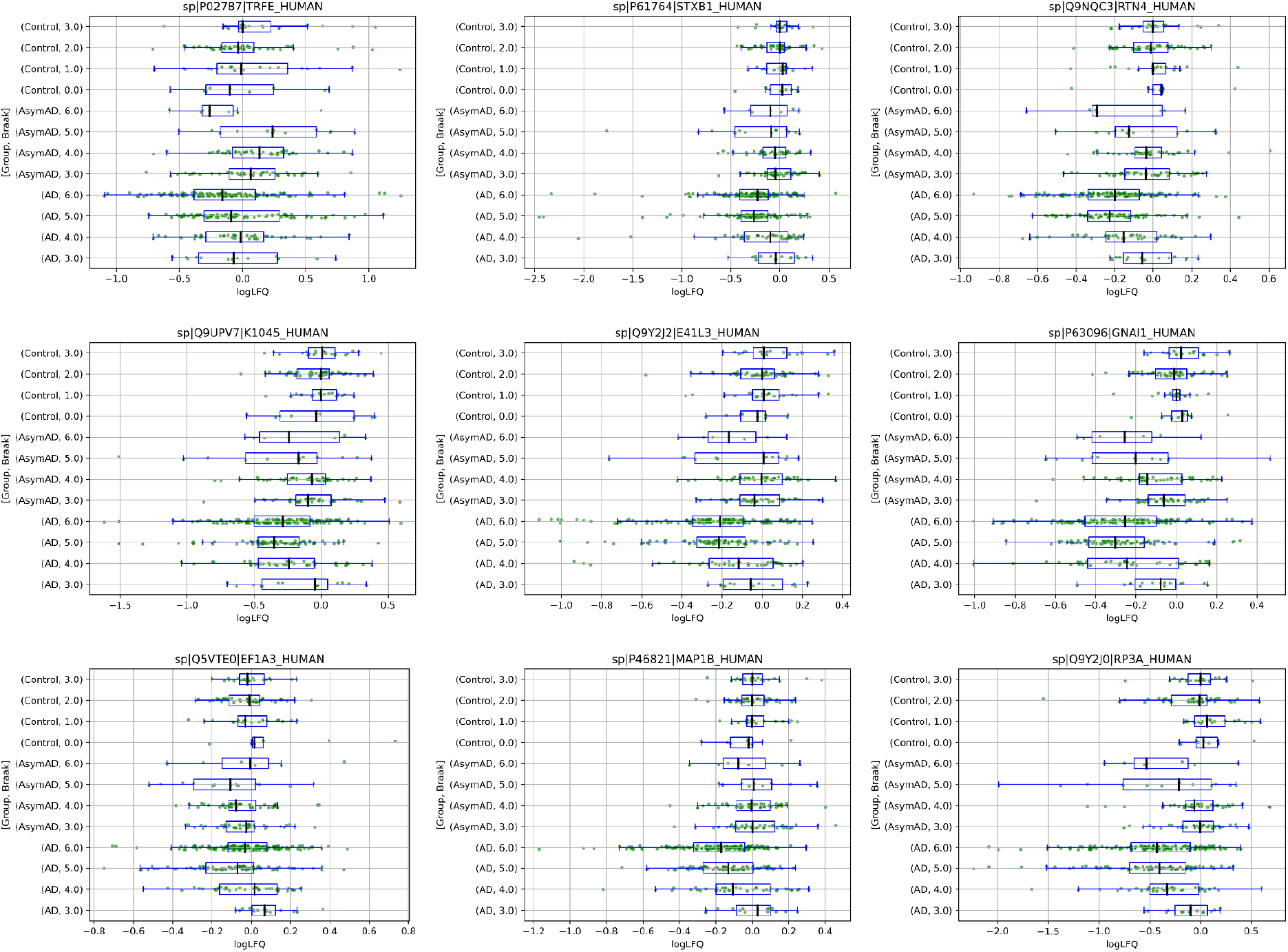
Label-free quantitation in log scale of proteins reported for RankedBraak and RankedCerad models grouped by Braak stage and condition.

## Results interpretation

Note that not all differentially expressed proteins detected in the Alzheimer group are really interesting. There are a lot of possible reasons for many proteins to be changed, sometimes significantly, as a consequence of cognitive issue development rather than being directly related. Table 4 shows all possible theoretical outcomes for protein expression for the study design used in this manuscript. Indeed, expression changes unique for the AD group may be explained by effects of patient treatment, issues with nutrition or hygiene, etc. Another ambiguous case is the protein expression change correlation between AsymAD and AD samples. Indeed, these changes are most likely a consequence of the presence of neurofibrillary tangles and neuritic plaques and do not provide much insight into the underlying biological mechanisms. Here, we focused on the protein expression changes unique for AsymAD group. After manual inspection of 27 proteins reported by RankedBraak, RankedCerad or AD_vs_AsymAD models (Figure 3 and Supplementary Figure S4), we found that only two proteins meet our criteria: TRFE (Serotransferrin) and APEX1 (DNA repair nuclease/redox regulator APEX1) (see Figure 4). APEX1 was not detected in standard differential expression analysis using DirectMS1Quant (Supplementary Table S1) and TRFE was detected only in two cases: AD_vs_AsymAD for ACT data set and AsymAD_vs_Control for BLSA data set.

**Table 4.** All possible outcomes for protein expression in different groups. 0 means no changes, + and - mean increase and decrease, respectively. Note that all protein LFQ values in our study were normalized to the Control group, which means only the “0” label could be used for the Control group.

| Group |  |  | Comment |
| --- | --- | --- | --- |
| Control | AsymAD | AD |  |
| 0 | 0 | 0 |  |
| 0 | 0 | + | Not interesting:<br>cannot be distinguished from treatment side-effects, nutrition problems, etc. |
| 0 | 0 | - |  |
| 0 | + | + | Not interesting:<br>does not help to find any mechanism unique for AsymAD group |
| 0 | - | - |  |
| 0 | + | 0 | Interesting: may explain why AsymAD group patients are cognitively normal during life |
| 0 | - | 0 |  |
| 0 | + | - |  |
| 0 | - | + |  |

**Figure 4.**
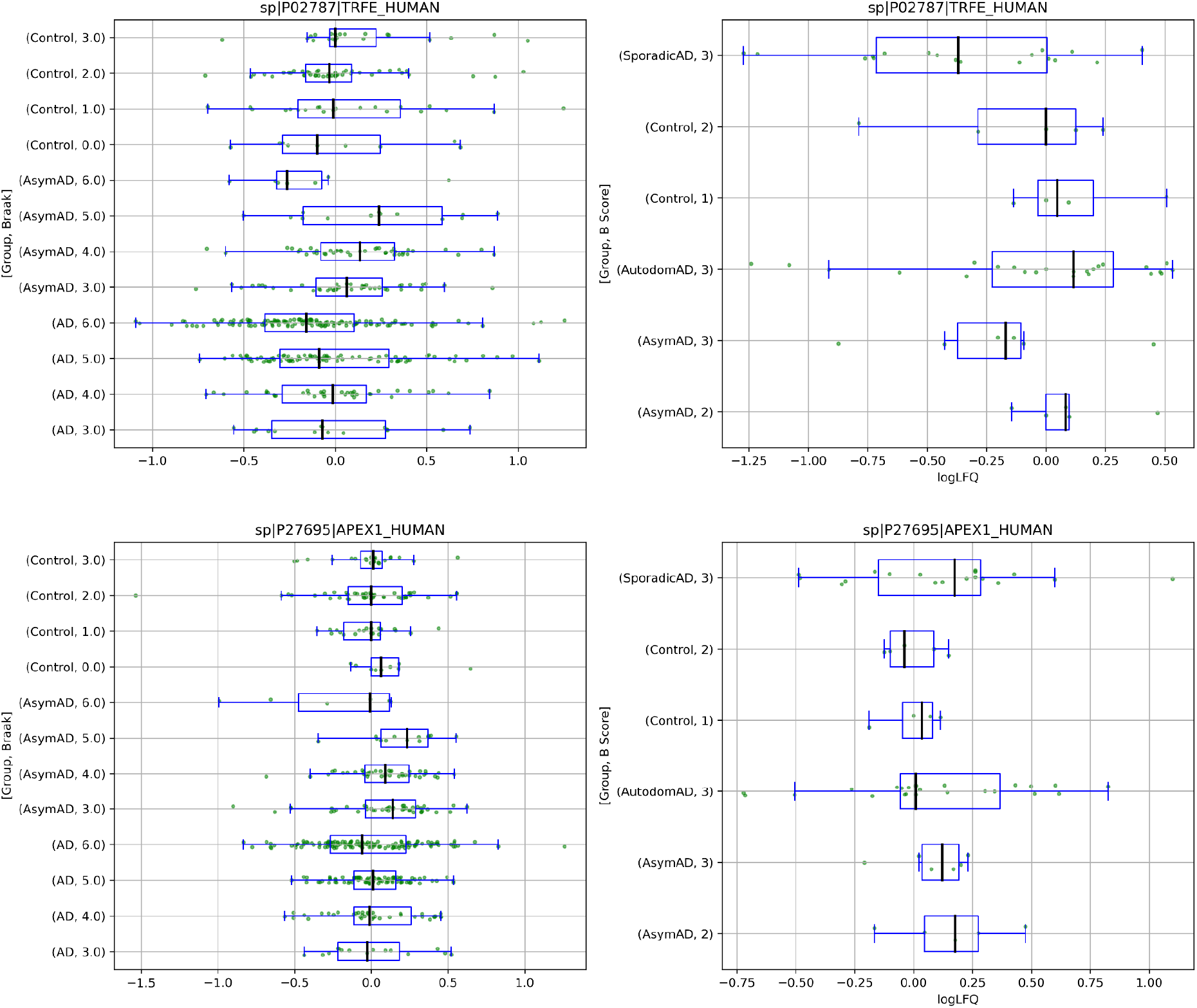
Label-free quantitation in log scale for TRFE and APEX1 proteins in DDA (left) and DIA (right) data grouped by Braak stage and condition. Note, that metainformation for DIA data contains B scores instead of Braak stages: B score 3 is equivalent to 5th and 6th Braak stages, 2 is 3rd and 4th stages, and 1 is 1st and 2nd stages.

APEX1 is a multifunctional protein that plays a central role in the cellular response to oxidative stress. Reportedly, APEX1 is a key inhibitor of cell death through ferroptosis.^15^ The latter is a recently proposed iron-dependent mechanism of cell death.^16,17^ It was noted that the level of iron significantly increases in the brain in case of cognitive dysfunctions.^18^ It was also shown that ferroptosis is accompanied with an increase in concentration of Fe^2+^ and Fe^3+^ ions in general and a decrease in the ratio of Fe^3+^/Fe^2+^.^19^ The latter study also showed 2.3- and 7.9-fold increase in Fe^2+^ and Fe^3+^ ion concentrations in mice brains with AD. The iron ions accumulate exclusively around Aβ amyloids which is explained by ion diffusion from surrounding cells. It was also shown that ferroptosis is a key mechanism of neurodegeneration in the familial form of AD in mice caused by mutations in presenilin 1 or 2.^20^ Finally, serotransferrin is responsible for the transport of iron ions as many studies suggest a correlation between its levels in plasma, serum and CSF with Alzheimer’s disease.^21–23^

To further elaborate on the ferroptosis’s AD’s hypothesis, we looked at the quantitation results for glutathione peroxidase 4 (GPX4), which is considered as a key ferroptosis inhibitor.^24^ Its concentration decreases for both AD and AsymAD groups with an increase in Braak stage (see Supplementary Figure S5), thus advocating for the ferroptosis being involved in the pathological process. Also note that GPX4 does not show any difference between AD and AsymAD groups, unlike the mentioned TRFE and APEX1. We hypothesize that the increase in TRFE and APEX1 in the AsymAD group represents a compensatory mechanism essential for neuronal survival in the face of accumulating misfolded proteins and the initiation of neurodegenerative processes. Supporting this hypothesis, aberrant iron distribution in the brain precedes tissue damage in both AsymAD and AD patients. This aberrant iron distribution, in turn, enhances the activity of ferroptosis as a cell death mechanism. Consequently, the increase in both TRFE and APEX1 likely acts as a key protective mechanism against the cell death in the AsymAD, potentially preventing the transition from the preclinical to clinical stages in Alzheimer’s disease pathogenesis. Thus, our findings demonstrate that ferroptosis takes place in brain tissue during the early stages of Alzheimer’s disease and that the regulatory mechanisms of ferroptosis may represent key events in the development of clinical forms. Therefore, early diagnosis and the development of novel targeted therapies involving ferroptosis may potentially extend the prodromal phase in AsymAD patients.

As a final comment, there are a few contradictory points in our findings. Firstly, the LFQ values for both TRFE and APEX1 in AsymAD group with Braak stage 6 are close to the ones in the AD group. In response, we may cautiously suggest that these samples may belong to the AD group but the patients died before exhibiting the cognitive dysfunction. Indeed, according to clinical guidelines^25^, Braak stages 5 and 6 are typically correlated with clinical dementia and are often classified as AD. Our observations for TRFE and APEX1 proteins advocate for the assignment of AsymAD with the Braak 6 stage to the AD group. However, the AsymAD group with the Braak 5 stage is more ambiguous and similar to the other AsymAD with Braak 3 and 4 stages. Secondly, the LFQ values grouped by data sets show that not all our findings are supported by the data sets studied, in particular, TRFE-AsymAD-MSSB, APEX1-AsymAD-BLSA, and APEX1-AD-DIA.

## Supporting Information

Supplementary information file includes upset plots for proteins reported as important by standard differential expression analysis and by machine learning approach; Label-free quantitation in log scale for different proteins grouped by data set and condition. (pdf)

Supplementary Table S1 contains the lists of differentially expressed proteins including their fold changes calculated by DirectMS1Quant for standard DE analysis. (xlsx)

## ACKNOWLEDGMENTS

This work, including development of the workflow based on the proposed modified machine learning algorithm, processing data sets, quantitative analysis of the proteomic data, as well as revealing differentially expressed proteins and their functionality was performed with financial support from the Russian Science Foundation, grant no. 23-45-00012. Jinghua Yang and Zhao Sun also thank the National Natural Science Foundation of China (project no. 82261138558) for supporting their work on selection and categorizing of label free proteomic data sets from publicly available repositories used in this study.

Authors thank Prof. Sergei Moshkovskii from Max Planck Institute for Multidisciplinary Sciences, Göttingen, Germany for useful discussion of the manuscript.

The results published here are in whole or in part based on data obtained from the AD Knowledge Portal (https://adknowledgeportal.org/).

ACT data set (syn5759376): These data were generated from postmortem brain tissue collected through the Adult Changes in Thought study (E. Larsen), and The University of Washington ADRC (T. Montine), with the proteomics provided by Drs. Seyfried, Lah, and Levey from Emory University (U01 AG046161; P50 AG025688; P30 NS055077).

MSSB data set (syn3159438): These data were provided by Dr. Levey from Emory University based on postmortem brain tissue collected through the Mount Sinai VA Medical Center Brain Bank provided by Dr. Eric Schadt from Mount Sinai School of Medicine.

Banner data set (syn7170616): These data were provided by Dr. Levey from Emory University. A portion of these data were generated from samples collected through the Sun Health Research Institute Brain and Body Donation Program of Sun City, Arizona. The Brain and Body Donation Program is supported by the National Institute of Neurological Disorders and Stroke (U24 NS072026 National Brain and Tissue Resource for Parkinson¹s Disease and Related Disorders), the National Institute on Aging (P30 AG19610 Arizona Alzheimer¹s Disease Core Center), the Arizona Department of Health Services (contract 211002, Arizona Alzheimer¹s Research Center), the Arizona Biomedical Research Commission (contracts 4001, 0011, 05-901 and 1001 to the Arizona Parkinson’s Disease Consortium) and the Michael J. Fox Foundation for Parkinson’s Research.

BLSA data set (syn3606086): These data were generated from postmortem brain tissue collected through The National Institute on Aging’s Baltimore Longitudinal Study of Aging and provided by Dr. Levey from Emory University.

