## Supplementary information file for "A modified decision tree improves generalization across multiple brains proteomic data sets and reveals the role of ferroptosis in Alzheimer’s disease"

**Table of Content**

**Figure S1.** UpSet plots for proteins reported as important by differential expression analysis and by machine learning approach. Page S-2.
**Figure S2.** Label-free quantitation of proteins for AD_vs_Control model (part 1/2). Page S-3.
**Figure S3.** Label-free quantitation of proteins for AD_vs_Control model (part 2/2). Page S-4.
**Figure S4.** Label-free quantitation of proteins for AD_vs_AsymAD and AsymAD_vs_Control models. Page S-5.
**Figure S5.** Label-free quantitation of GPX4 protein. Page S-6.

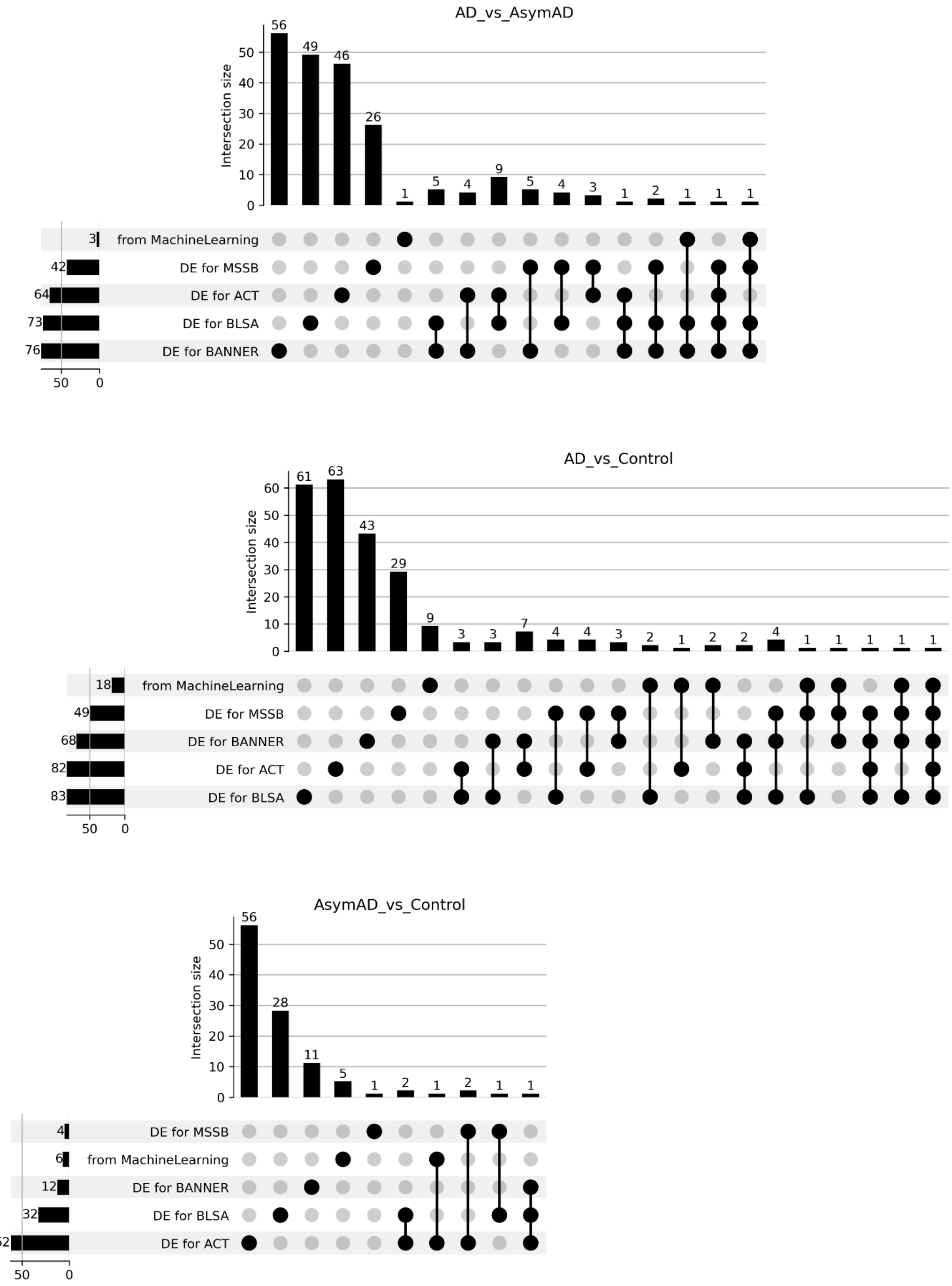


**Figure S1**. UpSet plots for proteins reported as important by either standard differential expression (DE) analysis in one of four data sets or by machine learning approach proposed in the study for (a) AD_vs_Control, (b) AsymAD_vs_Control and (c) AD_vs_AsymAD comparisons.

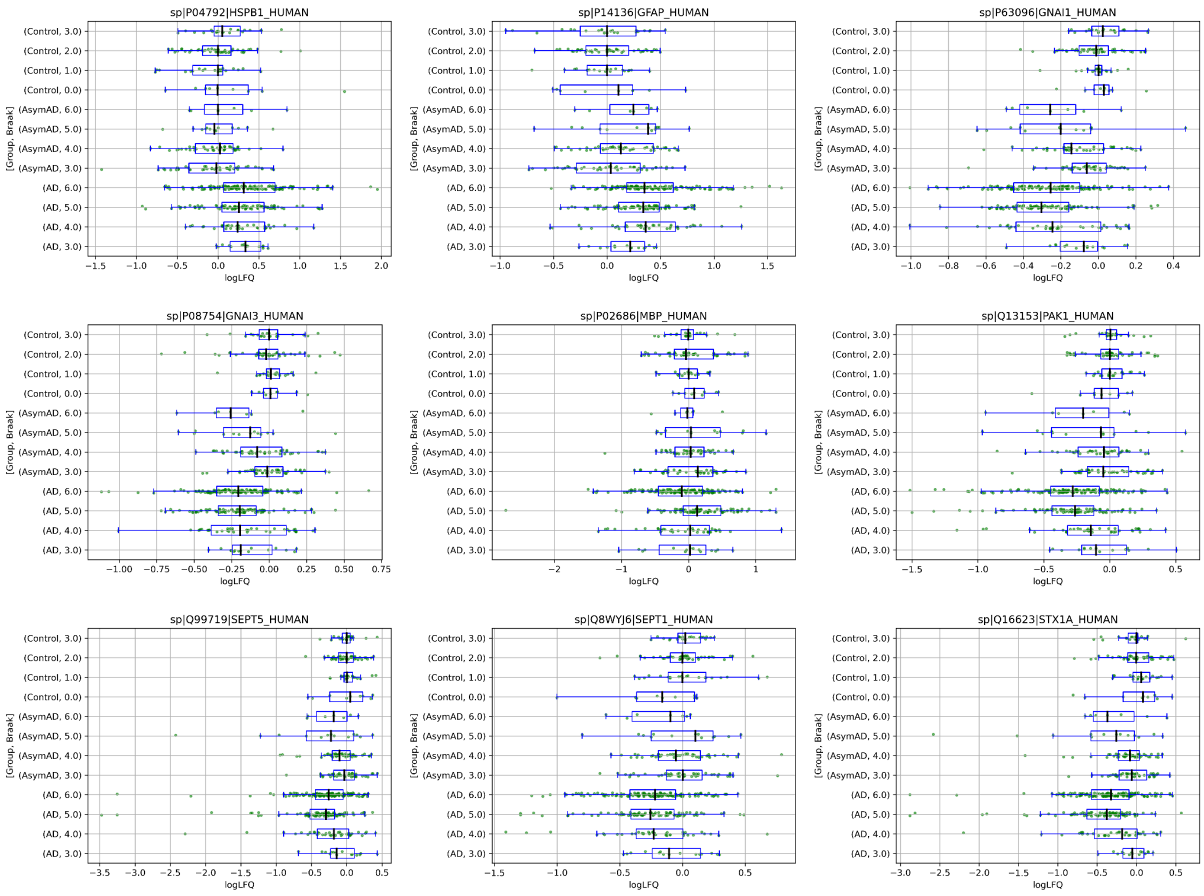


**Figure S2**. Label-free quantitation in log scale of ML-based reported proteins grouped by data set and condition for AD_vs_Control model (part 1/2).

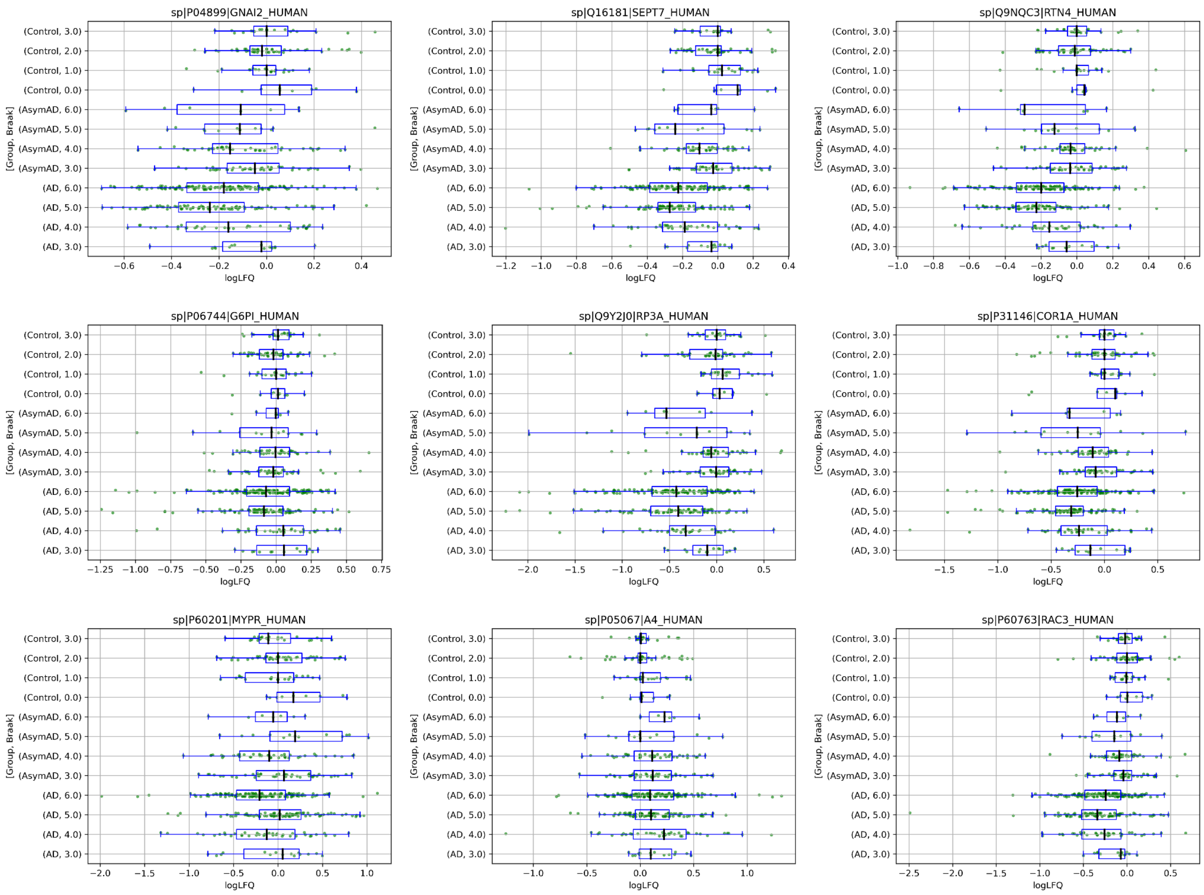


**Figure S3**. Label-free quantitation in log scale of ML-based reported proteins grouped by data set and condition for AD_vs_Control model (part 2/2).

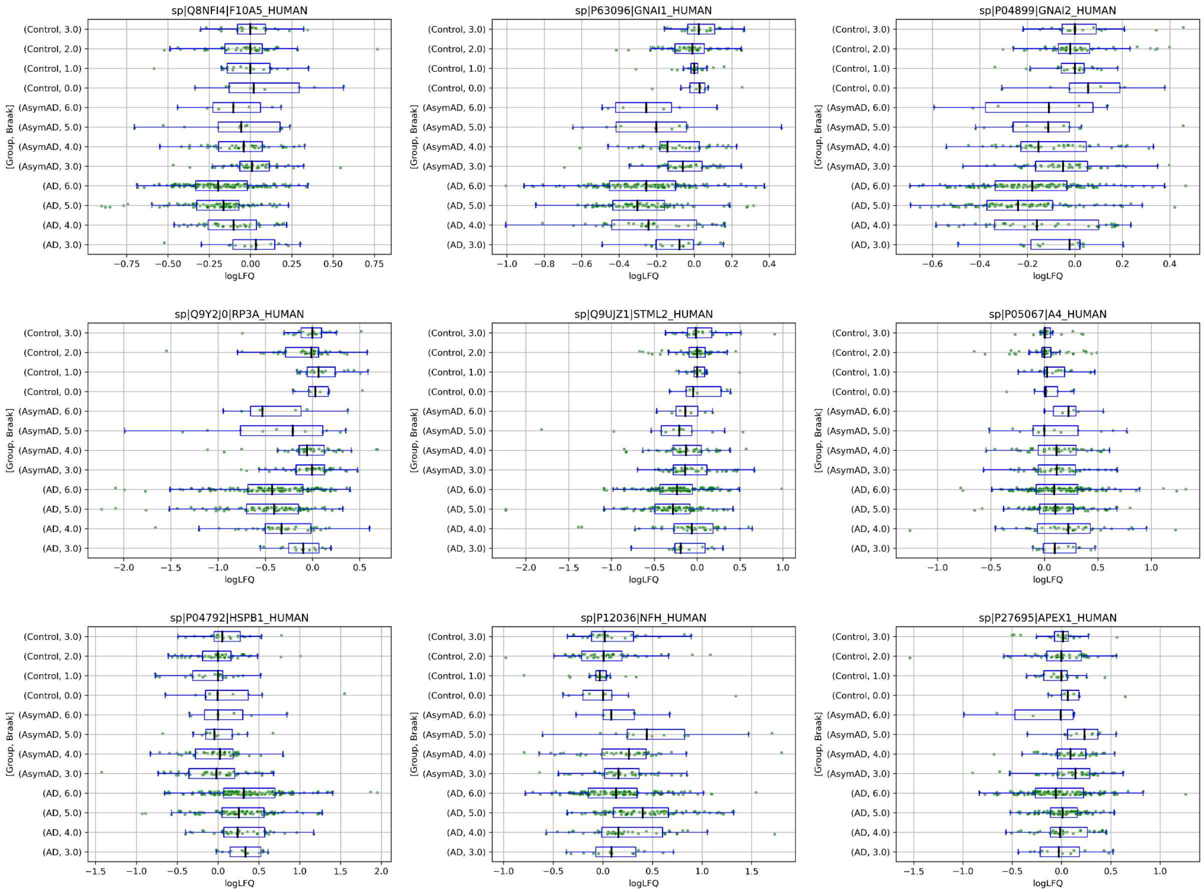


**Figure S4**. Label-free quantitation in log scale of ML-based reported proteins grouped by data set and condition for AsymAD_vs_Control and AD_vs_AsymAD models.

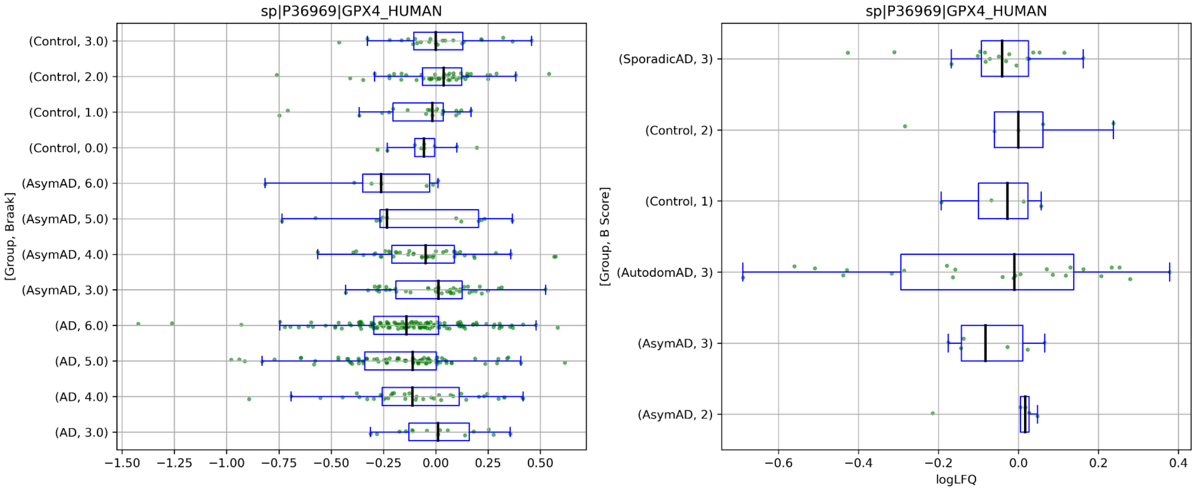


**Figure S5**. Label-free quantitation in log scale grouped by data set and condition for GPX4 protein .
